# Challenges in the Accurate Modelling of Lipid Dynamics in Monolayers and Bilayers

**DOI:** 10.1101/2024.09.12.612735

**Authors:** Carmelo Tempra, Víctor Cruces Chamorro, Titas Mandal, Salvatore Chiantia, Martin Vögele, Balázs Fábián, Matti Javanainen

**Affiliations:** Institute of Organic Chemistry and Biochemistry of the Czech Academy of Sciences, CZ-16000, Prague 6, Czech Republic; University of Potsdam, Institute of Biochemistry and Biology, DE-14476, Potsdam, Germany; Department of Theoretical Biophysics, MPI Biophysics, DE-60438, Frankfurt am Main, Germany; Unit of Physics, Tampere University, FI-33720 Tampere, Finland; Institute of Biotechnology, University of Helsinki, FI-00790 Helsinki, Finland

## Abstract

Recent advances in hydrodynamic theory have revealed the severe effect of periodic boundary conditions (PBCs) on the diffusive dynamics of lipid membranes in molecular dynamics simulations. Even when accounting for PBC effects, the corrected lipid diffusion coefficients often severely overshoot the experimental estimates. Here, we investigate the underlying reasons for the exaggerated dynamics, and suggest potential ways for improvement. To this end, we examine the diffusion of four lipid types in both bilayers and monolayers using the CHARMM36 force field. We account for PBC effects using the full hydrodynamic treatment: for bilayers we use non-equilibrium simulations to extract the interleaflet friction parameter used in the correction; whereas monolayer hydrodynamics are treated by setting this parameter to zero. Our results suggest that the dynamics of bilayers are too fast, even if interleaflet friction is accounted for. However, the change of the water model to OPC leads to an excellent agreement with experiments. For monolayers, the dynamics with the TIP3P water model agree well with experiments, whereas they are undershot with OPC. As OPC and TIP3P differ in both shear viscosity and surface tension, we develop two new mass-scaled water models to clarify the roles of the thermodynamic and kinetic properties of the water model on lipid dynamics. Our results indicate that both of these quantities play a major role in lipid dynamics. Moreover, it seems that the accurate description of diffusion in both lipid bilayers and monolayers cannot be accounted for by changes in the water model alone, but likely also requires modifications in the lipid model.

## 1 Introduction

All cells as well as the organelles in eukaryotic cells are encapsulated by compositionally complex, asymmetric, and heterogeneous membranes.^1–4^ The backbone of these membranes is provided by the lipid bilayer, whose structure has been carefully characterized over the past decades.^5^ The lipid bilayer consists of two monolayers facing each other such that their acyl chains form a hydrophobic core lined on both sides by the hydrophilic headgroups facing the aqueous solvent.

Because they are often easier to handle in experimental setups, lipid monolayers have been traditionally used as a proxy for bilayers^6^ but there are fundamental differences between the two lipid structures. Monolayers spread indefinitely at the air–water interface unless confined to a certain area per lipid (APL). In the laboratory setting, they are hence always studied under mechanical equilibrium, *i.e.* at a certain APL and the related surface tension *γ*. The APL–*γ* isotherms are often used to characterize the behavior of monolayers composed of lipids or other surfactants. In contrast, bilayers surrounded on both sides by a solvent have an equilibrium APL as the tensions of the two leaflets cancel out, ^7^ giving rise to structures such as vesicles. Thus, while bilayer phase behavior is only a function of the temperature at ambient pressure, the additional specification of APL (or *γ*) is required to fully characterize the state of a lipid monolayer. ^8^ Despite this fundamental difference, the lipids in the two arrangements are considered to be structurally similar, and Blume estimated that monolayers at a tension *γ* ≈ 30 mN/m would best correspond to bilayers.^9^

The acyl chains from opposing leaflets (*i.e.* monolayers) of a bilayer interact at the membrane core, which can strongly influence both lipid structure and dynamics. In some cases, acyl chains from one leaflet can penetrate into the opposing one, and this *lipid interdigitation* ^10^ can lead to composition-dependent structural perturbation. This penetration is not necessarily required for interaction, and the phase behavior of the two leaflets can be more generally affected by *interleaflet coupling* .^11–16^ Notably, significant theoretical effort has been invested into resolving the mechanisms that keep coexisting domains or phases in registry, that is, co-localized across the leaflets.^11,15–17^ The coupling effects are of particular interest in asymmetric systems ^13,14,18^ where phase separation only takes place in one leaflet. A prime example is the plasma membrane where only the extracellular leaflet is suggested display heterogeneity in the form of ordered domains.^19^

While interleaflet coupling is somewhat of an ambiguous concept, the *interleaflet friction b* captures these complex interactions in form of a single well-defined parameter. It describes the speed at which the leaflets of a bilayer drift with respect to each other under shear forces (see section “Experimental”). This parameter is of special interest in hydrodynamic models of lateral diffusion. While the celebrated Saffmann–Delbrück model^20^ links diffusion coefficients of trans-membrane objects to the properties of the diffusing object and the host membrane, *b* needs to be included in the model to accurately capture the movement of monotopic objects such as lipids. ^21–24^ Still, relatively few values of *b* have been reported in the literature, and these values are surprisingly scattered, especially across different measurement techniques.^25^

Thanks to the constant improvement in the accuracy of force fields and the available spatial and temporal scales, molecular dynamics (MD) simulations have become a standard tool in the study of biomembranes.^26–28^ As the different experimental membrane models— *e.g.*, supported membrane, black lipid membrane, free-standing membrane, or a monolayer in a Langmuir trough—have their strengths and weaknesses,^6,29^ so do the measurement techniques. MD simulations allow the study of monolayers and bilayers under similar conditions and without the need for complicated labeling techniques such as, *e.g.*, fluorescent probes. Thus, they are an excellent choice of method to compare their behavior.

Although common atomistic simulation models represent the structures of lipid bilayers reasonably well,^30–32^ they struggle in capturing the dynamic properties—reorientation of lipid segments^33^ and the diffusion of lipids along the membrane plane. ^32^ Notably, simulations are essentially always performed using periodic boundary conditions (PBCs), which lead to correlations in lipid motion due to the long-ranged nature of hydrodynamic flow fields.^34^ Fortunately, we can account for these effects in lipid membranes,^24,35,36^ which allows the direct comparison between simulation and experiment. While the first systematic studies comparing the lateral diffusion coefficients from MD simulations and experiments have emerged,^32,35,37,38^ no such comparison of diffusion in monolayers has been performed to our knowledge. Regarding interleaflet friction, there are multiple strategies available for extracting it from non-equilibrium^39–42^ or equilibrium^43^ simulations. Most of the studies are performed at a coarse-grained resolution for which the potential energy surface is smoother than at atomistic resolution and the extracted *b* values are an order of magnitude smaller than those from state-of-the-art tank-treading experiments.^25^ Moreover, the values of *b* have, to our knowledge, not been explicitly measured in specific membranes and then subsequently used as an input parameter when accounting for PBC effects in the same membranes.

We thus identified the following open questions regarding the dynamics of lipid monolayers and bilayers: 1) Do atomistic MD simulations capture the lateral diffusion coefficients in monolayers and bilayers when interleaflet friction is properly accounted for, or do other parameters such as the non-zero surface tension of monolayers play a role?; 2) Do atomistic MD simulations capture the interleaflet friction values from latest tank-treading experiments?; and 3) If the answers to the previous two questions are negative, what methodological developments are required to obtain better agreement with experiment? To answer these questions, we performed atomistic MD simulations of four single-component monolayers and bilayers with the CHARMM36 force field. Our simulations highlight how the overly large diffusion coefficients, their box-size dependencies, and additional effects in monolayers due to the use of inaccurate water models obfuscate the comparison of MD simulations and experiments. We demonstrate that adapting the water model to better reproduce experimental properties largely cures the issues present in bilayer simulations. However, the situation with monolayers is more complicated, and might also require changes in the lipid model.

## 2 Results and Discussion

We simulated four single-component lipid bilayers (Fig. 1B) and the corresponding monolayers (Fig. 1C), each in three different sizes. The four lipid types, each with one oleate chain, are shown in Fig. 1A. The APL in the monolayer was set to match that of the relaxed lipid bilayers. All 24 simulations (4 lipid types, 3 sizes, monolayer & bilayer) were run for 1 µs using the GROMACS simulation engine ^44^ and the CHARMM36 lipid force field^45^ together with the CHARMM-specific TIP3P water.^46,47^ The simulation data for bilayers (10.5281/zenodo.7103807) and monolayers (DOI: 10.5281/zenodo.11219521) are available in the Zenodo repository. The simulations with the largest system size are visualized in a movie available at DOI:10.6084/m9.figshare.21187600.v1.

**Figure 1:**
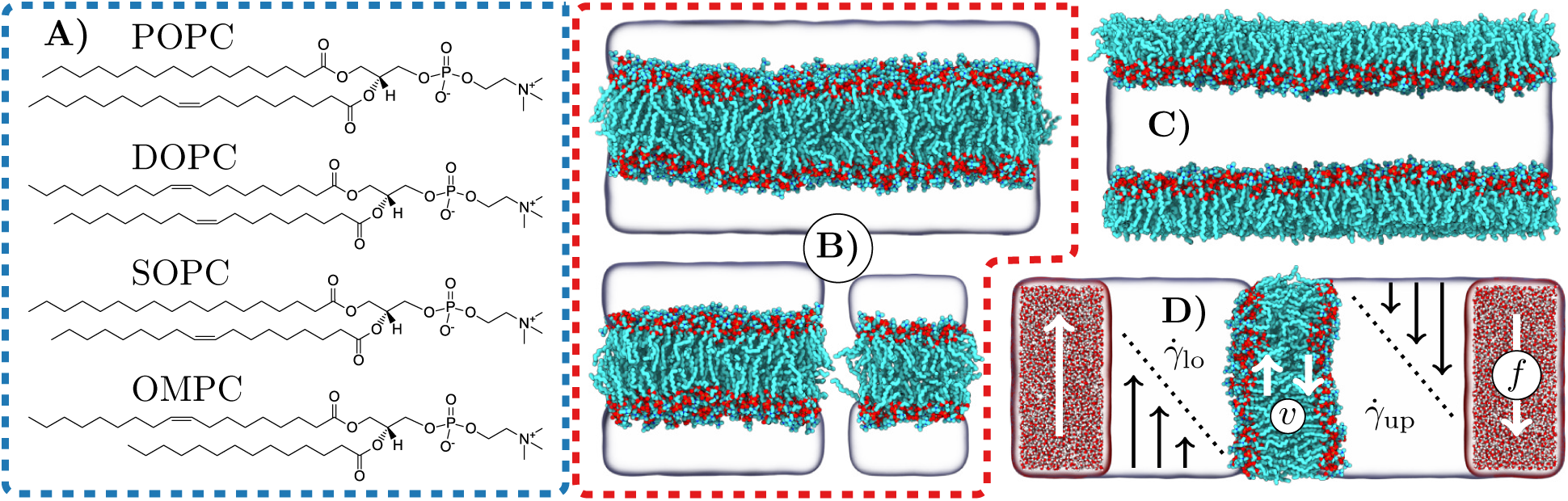
Summary of simulated systems. A) Chemical structures of the studied lipids. B) Example snapshots of the POPC bilayers after 1 µs of simulation. The three system sizes are shown; large (top), medium (bottom left), and small (bottom right). Lipids are coloured based on chemical elements (cyan: carbon, red: oxygen, blue: nitrogen, and brown: phosphorus). Water is shown as a transparent surface and hydrogens are omitted. C) A snapshot of the setup containing two large POPC monolayers after 1 µs of simulation. Colouring as in B). The vacuum slab separating the lipid acyl chains across the periodic boundary conditions is omitted. D) A snapshot and a schematic of the shearing simulations. Compared to B), the thickness of the water slab is increased. Opposite forces (*f* ) are applied to the regions highlighted in red and with explicit water molecules rendered. The shearing gives rise linear velocity profiles (*ζ̇*) in the water near the membrane surface, which further induces a relative drift (*v*) of the membrane leaflets. Colouring as in B).

We first confirmed that the equal APL of the corresponding lipid bilayers and monolayers leads to similar structures sampled by the lipid molecules. Our recent work demonstrated the similarity of the density profiles of the monolayer and a single bilayer leaflet, indicating that the overall dimensions such as thickness are very similar. ^48^ In addition, we calculated deuterium order parameter profiles (−*S*_CD_) along the lipid acyl chains, which describe the average lipid conformations (see section “Experimental”). These profiles, shown in Fig. S2 in the Supplementary Information (SI), indicate that an equal APL ensures that the individual lipids also sample similar conformations in monolayers and bilayers. Bilayers at their equilibrium APL (values listed in Table 1) are overall tensionless, but monolayers at the air–water interface are always under tension. The surface tension values (*γ*) of the lipid covered interfaces are listed in Table 1, and they are generally *γ* ≈30 mN/m. As the surface pressure is defined as the difference of surface tension between the pure and lipid-covered interfaces (Π = *γ*_0_ − *γ*) and (CHARMM-specific) TIP3P water having *γ*_0_ ≈ 50 mN/m,^49^ the surface pressures of the studied lipid systems monolayers are Π ≈ 20 mN/m. This agrees well with experimental predictions.^9^

**Table 1:** Properties of the bilayers and monolayers from equilibrium simulations. The APL and *γ* values are mean ± standard deviation of the values for the three system sizes. The diffusion coefficients are the values for an infinite system, *i.e.* with the PBC effects eliminated. Diffusion coefficients are extracted for bilayers using the *b* values obtained from non-equilibrium shearing simulations 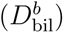. Comparison to values extracted using the numeric approach ^36^ or the analytical approximation (Eq. (1)), both with the assumption of an infinite interleaflet friction, are shown in Table S5 in the SI), and they differ by up to 15%. For monolayers, we used 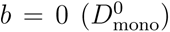. The individual values for all system sizes are shown in Tables S1 (TIP3P), S2 (OPC), and S3 (TIP3P^S^ & OPC^S^) in the SI. All these values, including the 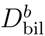 are also shown in Fig. S1 in the SI. Units are as follows: APL in Å^2^, *b* in MPa·s/m, *D*s in 10^−8^ cm^2^/s, *µ* in mPa·s, *L*_SD_ in nm, and *γ* in mN/m. *^∗^*) The interleaflet friction for the POPC bilayer with other water models was assumed to be the same as with TIP3P water.

|  | Bilayer |  |  |  |  | Monolayer |  |  |  |
| --- | --- | --- | --- | --- | --- | --- | --- | --- | --- |
| Lipid | APL | $D_{\text{bil}}^b$ | $b$ | $\mu_{\text{bil}}$ | $L_{\text{SD}}^{\text{bil}}$ | $\gamma$ (mN/m) | $D_{\text{mono}}^0$ | $\mu_{\text{mono}}$ | $L_{\text{SD}}^{\text{mono}}$ |
| TIP3P water model |  |  |  |  |  |  |  |  |  |
| OMPC | $64.5 \pm 0.1$ | 15.4 | $32.3 \pm 2.8$ | 47.3 | 276 | $25.5 \pm 2.5$ | 34.8 | 32.4 | 189 |
| DOPC | $67.7 \pm 0.0$ | 13.1 | $46.6 \pm 2.0$ | 65.5 | 392 | $30.1 \pm 1.8$ | 33.8 | 32.8 | 197 |
| POPC | $63.7 \pm 0.1$ | 13.2 | $30.1 \pm 3.6$ | 56.4 | 338 | $31.8 \pm 1.4$ | 29.3 | 41.4 | 248 |
| SOPC | $63.4 \pm 0.1$ | 12.3 | $33.4 \pm 4.7$ | 57.4 | 344 | $31.7 \pm 1.6$ | 22.9 | 58.5 | 351 |
| OPC water model |  |  |  |  |  |  |  |  |  |
| OMPC | $63.3 \pm 0.2$ | 7.2 | $32.3 \pm 2.8^*$ | 100.3 | 233 | $28.3 \pm 4.4$ | 11.6 | 90.5 | 279 |
| DOPC | $66.0 \pm 0.1$ | 6.6 | $46.6 \pm 2.0^*$ | 117.2 | 286 | $29.9 \pm 1.7$ | 13.7 | 81.4 | 199 |
| POPC | $62.4 \pm 0.1$ | 5.8 | $30.1 \pm 3.6^*$ | 135.0 | 342 | $29.3 \pm 1.1$ | 11.7 | 94.8 | 240 |
| SOPC | $62.2 \pm 0.2$ | 5.3 | $33.4 \pm 4.7^*$ | 146.1 | 376 | $29.5 \pm 1.2$ | 10.7 | 101.4 | 261 |
| Model | POPC with mass-scaled water models |  |  |  |  |  |  |  |  |
| TIP3P<sup>S</sup> | $64.0 \pm 0.1$ | 11.3 | $30.1 \pm 3.6^*$ | 61.1 | 219 | $26.8 \pm 3.1$ | 19.2 | 67.5 | 242 |
| OPC<sup>S</sup> | $62.5 \pm 0.1$ | 6.7 | $30.1 \pm 3.6^*$ | 48.7 | 445 | $31.1 \pm 1.3$ | 14.3 | 32.8 | 301 |

Having established the structural similarity of monolayers and bilayers, we proceeded to study lipid diffusion. We extracted the diffusion coefficients of the lipids with state-of-the art tools, properly accounting for fluctuations in simulation system size due to pressure coupling (bilayer simulations), and providing well-defined error estimates.^38,50^ The diffusion coefficient values extracted for monolayers and bilayers, all lipid types, and all system sizes are listed in Table S1 in the SI. These values themselves cannot be compared to experiments, as the use of periodic boundary conditions results in correlations of lipid motion. ^23,24,34–36^ To correct for the periodic boundary conditions, we extrapolated the diffusion coefficients to the infinite system size using Eq. (1).^36^ To perform the correction, we also need information on other system properties, such as simulation box dimensions, membrane thickness, and the shear viscosity of water. Moreover, the monolayers and bilayers require different treatments. In our monolayer simulations, we effectively set up the two leaflets of a lipid bilayer, but separated by a large slab of vacuum (≈17.5 nm, omitted from Fig. 1C). Hence, we model our monolayer as a bilayer with a vanishing interleaflet friction coefficient *b* = 0 and with the same membrane thickness as calculated for the corresponding bilayer.

In bilayers, the interleaflet friction contributes to the dynamics, and we handled it in two ways: We first assumed infinite friction *b* = ∞, which effectively corresponds to two lipids in the opposing leaflets diffusing as one membrane-spanning entity. In addition, we also used the *b* values extracted from shearing simulations (see Fig. 1D). In these simulations, we applied forces in opposite directions (*f* ) to slabs of water far away from the membrane, resulting in a velocity gradient in the thick water slab and hence a shear rate (denoted by *ζ̇* instead of the commonly used *γ̇* to avoid confusion with the surface tension) in both membrane leaflets. The total shear rate (*ζ̇*_total_ = *ζ̇*_upper_ + *ζ̇*_lower_) induced a displacement of the leaflets with respect to one another at a constant velocity *v*. We calculated *b* from the relations of the shear force; *F* = *µ_w_ζ̇*_total_ and *F* = *bv* (see section “Experimental” for details). Overall, the velocity gradient of the non-sheared water regions depended linearly on the shearing forces applied to water molecules distant from the bilayer (Fig. S10A), and the drift of the bilayer leaflets also depended linearly on this gradient (Fig. S10B). This indicates that the applied forces were suitably large to induce an effect in the bilayer, yet did not result in turbulent effects or structural alterations therein. The interleaflet friction values obtained from shearing simulations are shown in Fig. 2 and also listed in Table 1. All the parameters from the shear simulations, including the values of *b* from the simulations with different shear rates, are available in Table S4 in the SI.

**Figure 2:**
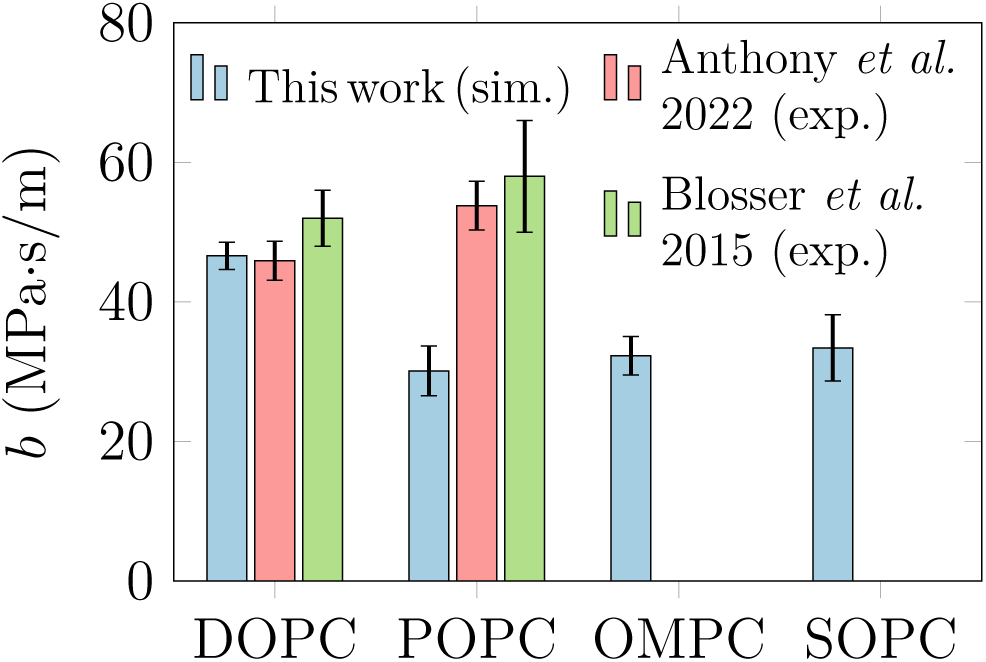
Interleaflet friction from non-equilibrium shearing simulations. The values from simulation are extracted using five different shearing rates, and the mean ± standard deviation of these values are shown. The independent shear rates are shown in Fig. S10. The experimental values, measured using shear-driven tank treading of supported lipid bilayers, are taken from Refs. 25 (Anthony, 2022) and 51 (Blosser, 2015).

Overall, we find DOPC to have the highest *b* ≈ 50 MPa·s/m, and this behavior is reproduced across the five studied shearing rates (Fig. S10C). Other studied lipids show mutually similar values of *b* ≈ 30 MPa·s/m, again consistent across various shearing rates. Experimental values obtained using different techniques span a large range, yet shear-driven supported lipid bilayer tank-treading experiments provide consistent values for lipids,^25,51^ and values for DOPC and POPC from such experiments are also shown in Fig. 2. For DOPC, the agreement seems to be excellent, whereas for POPC our simulations provide a smaller estimate than the experiments. However, an earlier simulation study with the same CHARMM36 lipid force field found a somewhat smaller value of *b* ≈ 20.9 ± 0.1 MPa·/ms for DOPC.^39^ The origin of this discrepancy is unclear; our higher value is consistent between the five different shearing rates (Fig. S10C, yet the value from Ref. 39 would better fit the trend observed in our simulations. Moreover, our structural analyses of the lipid bilayers^48^ revealed similar localization of the acyl chain termini for both POPC and DOPC, hinting at similar ruggedness at the molecular scale. As we demonstrate below, a difference in *b* at this level (≈50 vs. ≈20 MPa·s/m) leads to a 5% difference in the PBC-corrected diffusion coefficients, and hence does not affect any of our conclusions.

With the values of *b* established, we proceeded to extract the lateral diffusion coefficients, *D*, from the monolayer and bilayer simulations. We obtained them by fitting the diffusion coefficients from our simulations at various box widths to the finite-size correction, ^36^ using the membrane surface viscosity *η_m_* as the free parameter and the value of *b* from our shearing simulations. In Table S5, we show how sensitive the *D* values are to the used fit function: We compare values obtained using the full numerical treatment for monotopic inclusions as done here in the main text 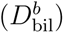 to a similar treatment assuming lipids to be bitopic inclusions 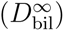. In the latter, two lipids are considered to diffuse as an interleaflet dimer, corresponding to *b* = ∞. We also included the values obtained using an analytical approximation for bitopic inclusions, Eq. (1) 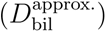. For the monolayers, *b* = 0 was always used, resulting 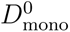. The 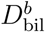 and 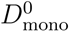 values are compared to experimental values extracted in our previous work ^48^ in Fig. 3.

**Figure 3:**
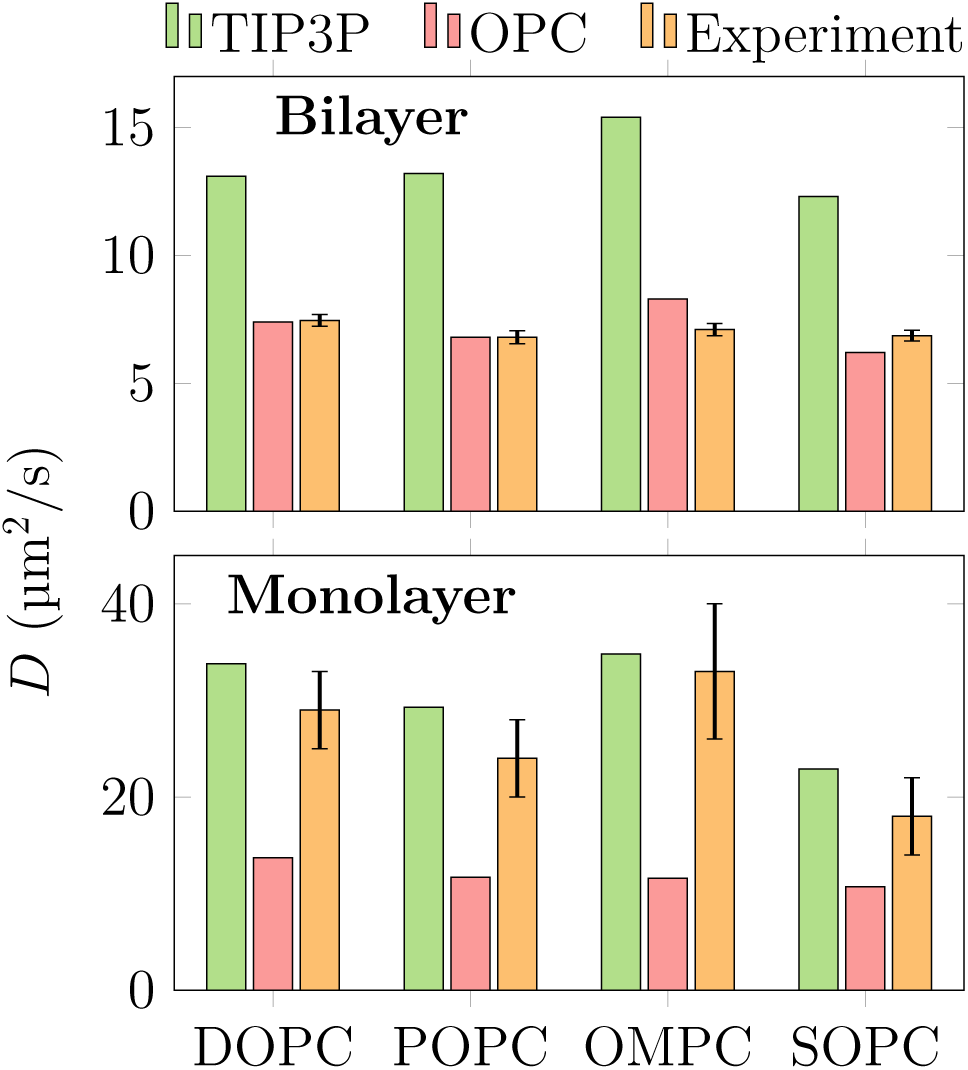
Diffusion coefficients from simulations and experiments. The diffusion coefficient values from simulations have been corrected for PBC effects. The experimental values are taken from our recent work.^48^

The diffusion coefficients obtained from simulations of bilayers (see “TIP3P”) overshoot the experimental values by a factor of ≈2. To eliminate the possibility that the experiments ^48^ are affected by the presence of fluorescence probes, we compared also to earlier label-free pulsed field gradient NMR measurements. Filippov et al. found values of 8.3 µm^2^/s and 7.8 µm^2^/s for DOPC and POPC at 298 K,^52^ in good agreement with the experiments shown in Fig. 3. These results confirm previous studies ^32,37^ which found that diffusive dynamics in CHARMM36 bilayers are significantly overestimated. We reproduce this finding even when interleaflet coupling is explicitly included in the PBC correction. Importantly, the value obtained using an interleaflet friction coefficient of 20.9 MPa·/ms^39^ in the fit, we obtained a value 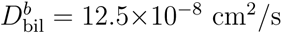, *i.e.* a 5% lower value than the one reported in Table 1.

Even worse, the simulations do not even capture the trend between the different lipid types. Experimentally, DOPC*>*OMPC*>*SOPC*>*POPC, whereas the simulations report OMPC*>*POPC*>*DOPC*>*SOPC. Hence, the mutual order between POPC and DOPC in the simulations also disagrees with the aforementioned label-free NMR experiments.^52^ Notably, the simulated trend is more complex than the simple free area theory, ^53^ as the diffusion coefficients do not show an exponential dependence on APL (Fig. S5 in the SI). Still, the differences, especially among experimental values, fall within the error estimates, and estimating the error in the simulation values is challenging due to the fitting approach used to eliminate finite size effects (Eq. (1) in section “Experimental”). Errors are available for values of the individual simulations, yet even their relative values cannot be compared to experiments due to the different magnitudes of the corrections for each lipid type.

The diffusion coefficients of the simulated monolayers, 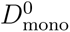, agree very well with the experimental ones both in terms of the absolute values as well as the trend between lipid types (see “TIP3P” in Fig. 3). The values from simulations are overall twice as large as for the bilayers (Fig. S4 in the SI), whereas on the experimental side, the corresponding speedup is 4-fold (Fig. 3). Indeed, the monolayer simulation reproduces the experimental trend (OMPC*>*DOPC*>*POPC*>*SOPC) and—unlike in the case of bilayers—the differences between lipid types are significantly larger. The error estimates are also larger, but apart from OMPC and DOPC (*p* = 0.18), the differences are significant (*p* = 0.01 for DOPC and POPC and *p <* 0.001 for other pairs).

It might at first seem that CHARMM36 is able to capture the lateral dynamics in monolayers but the comparison with bilayers reveals a more complicated picture. Monolayers and bilayers differ by the presence of interleaflet coupling, so it is suspicious that—at least in the case of DOPC—both monolayer diffusion coefficient and interleaflet friction agree with experimental values, while the bilayer diffusion coefficient does not. The reference experiments used the same fluorescent probe for both monolayers and bilayers,^48^ and hence its effect can be ruled out. One well-known limitation of biomolecular force fields, including CHARMM36, is that their recommended standard TIP3P water model severely underestimates the experimental surface tension. At 298 K, the experimental value is *γ*_0_ ≈72 mN/m, whereas that of TIP3P is *γ*_0_ ≈50 mN/m.^49^ Therefore, it is possible that the good agreement of the diffusion coefficients between our monolayer simulations and experiments could result from a favorable cancellation of errors; the dynamics of the lipids are inherently too fast as evidenced by the bilayer simulations, but the incorrect surface tension and the resulting incorrect surface pressure of the monolayer could slow diffusion down.

We examine the influence of surface tension on the lateral diffusion coefficient by changing the water model in our simulations. We repeated all 24 simulations with the recent “Optimal Point Charge” (OPC) water model,^54^ which reproduces the experimental surface tension of neat water.^49,55^ These simulation data are also available in Zenodo under DOIs: 10.5281/zenodo.11220606 (POPC) and 10.5281/zenodo.13355257 (other lipids). The bilayer areas per lipid, monolayer surface tensions, and the diffusion coefficients for each lipid and each system size with OPC water are listed in Table S2, while the averages over three system sizes are listed in Table 1. The change of the water model from TIP3P to OPC was shown to have a relatively small effect on the structural properties of tensionless bilayers.^55^ Contrarily, the choice of water model significantly affects the behavior of monolayers, especially at larger APLs, where the smaller cost of exposing the water surface leads to the opening of pores in the monolayer.^55^ The monolayer APLs studied here—matching those of the bilayers—are high, and are barely affected by the change of the water model (Fig. S3). Still, the change of the water model also leads to a slight condensation of the bilayers, as the APL decreases by a few percent upon the change from TIP3P to OPC (see Tables 1 as well as S1, S2, and S3 in the SI).

The change in water model leaves the surface tension of the monolayers unchanged, it must therefore increase the surface pressure. The surface tension of OPC is ≈20 mN/m larger than that of TIP3P. Hence, a simple assumption would be that if monolayer surface pressure remains constant, the tension of the lipid-covered interface increases by this ≈20 mN/m upon changing the water model. However, the surface tension of the simulated OPC monolayers is actually 29.3 ± 0.6 mN/m (mean and standard deviation over the lipids), *i.e.* essentially identical to the 29.8 ± 2.6 measured in TIP3P monolayers (see Table 1). These results indicate that using the OPC water model with correct surface tension increases the surface pressure of the monolayers by ≈20 mN/m to ≈40 mN/m, counteracting the change in *γ*_0_.

We now investigated how this change in the surface pressure affect the lateral diffusion coefficient in the monolayers despite the constant APL. We repeated the diffusion coefficient calculation for the bilayer and monolayers with OPC water, and used the corresponding shear viscosity, *µ*_w_,^56^ for the PBC corrections. The corrected values are listed in Table 1 and shown in Fig. 3.

Our results indicate that OPC with a higher shear viscosity (*µ*_w_ ≈ 0.80 mPa·s^56^ versus *µ*_w_ ≈ 0.32 mPa·s calculated for TIP3P^57^) significantly slows down lipid diffusion in both mono- and bilayers. In bilayers, this slowdown of more than 50% leads to the simulations actually being in good agreement with the experimental estimates. Hence, the too rapid diffusive dynamics reported earlier for CHARMM36^32,37^ seems to be largely due to the insufficient shear viscosity of the TIP3P water model.

However, the situation with the monolayers is less clear, as the change from TIP3P to OPC not only increases the shear viscosity by a factor of ≈2.5, but the different surface tension also plays a role in the monolayer setup. Overall, the change of water model slows down diffusion in monolayers by over 50%, similar to the corresponding effect in bilayers. Thus, viscosity might play a significant role, whereas the surface tension might not. This is indeed supported by the fact that the surface tension of the lipid-covered air–water interface is essentially unchanged upon the change of the water model.

Unfortunately, it is challenging to separate the contribution of the surface tension from that of the shear viscosity. As shown in Fig. S7 in the SI, these two properties show strong correlation between the available water models. We also looked into a recently parametrized set of water models, in which density, self-diffusion coefficient, the first peak in the oxygen– oxygen radial distribution function, and the dielectric constant were used as target properties.^58^ Unfortunately, this set contained no models with poor surface tension or poor shear viscosity values to help us eliminate one of the two variables (Fig. S7 in the SI).

As developing models with good (bad) surface tension and bad (good) shear viscosity seemed unfeasible, we decided to develop two water models whose viscosities were similar (although not correct), whereas the surface tensions were very different—one agreeing with the experiment. The best way to proceed is to transform the TIP3P and OPC water models alchemically by increasing the atom masses by a ratio of *M* . Scaling the masses changes the shear viscosity as 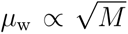, without altering any thermodynamic properties such as the surface tension.^59^ Thus, we developed a pair of new OPC- and TIP3P-based water molecules, coined OPC^S^ and TIP3P^S^, with equal (but incorrect) viscosities, without altering their thermodynamic properties such as surface tension. For shear viscosity, we indeed observed the expected scaling with respect to mass (Fig. S8 in the SI), and an equal shear viscosity value of *µ*_w_ ≈ 0.54 mPa·s was found when the masses of TIP3P and OPC were scaled by factors of 2.9 and 0.4, respectively. We also confirmed that the change in mass did not indeed affect surface tension (Fig. S8 in the SI). We then repeated the POPC bilayer and monolayer simulations with the modified TIP3P^S^ and OPC^S^ water models.

The diffusion coefficients of POPC bilayers and monolayers with the standard and mass-scaled water models are shown in Fig. 4A & B. Importantly, the corresponding water models should provide identical thermodynamic yet different dynamic behavior, whereas both mass-scaled models should provide identical dynamic yet different thermodynamic behavior. Unsurprisingly, an increase (decrease) of shear viscosity upon mass-scaling the TIP3P (OPC) water model leads to a decrease (increase) of the the diffusion of lipids in a bilayer. Curiously, the mass-scaled models—despite having similar viscosities—still provide very different lipid diffusion coefficients, indicating that their thermodynamic properties also play a significant role. Therefore, it seems likely that increasing the shear viscosity of water, *µ*_w_, alone is not adequate to cure the overly fast dynamics of lipids in bilayer simulations with standard TIP3P. Regarding the diffusion coefficients in monolayers, we observe the same trend as for the bilayer with the different water models. However, both mass-scaled models undershoot the experimental value, and only the original TIP3P—with its poor shear viscosity and surface tension—is able to match it. The deviation from experimental estimates for monolayers and bilayers is shown in Fig. 4C, and demonstrates that capturing the experimental values is infeasible by simply changing the water model. As a further illustration, Fig. 4D highlights the deviations (marker size) for bilayers and monolayers as a function of the two key water parameters—shear viscosity and surface tension. Based on this plot, one could speculate that a model with a shear viscosity of ≈60 mPa·s and a surface tension of ≈60 mN/m could provide the smallest least-squares deviation from experimental values. Hence, in studies where the reproduction of dynamics in both bilayers and monolayers is essential, TIP4P and SPC/E should be considered as potential water model candidates based on Fig. S7.

**Figure 4:**
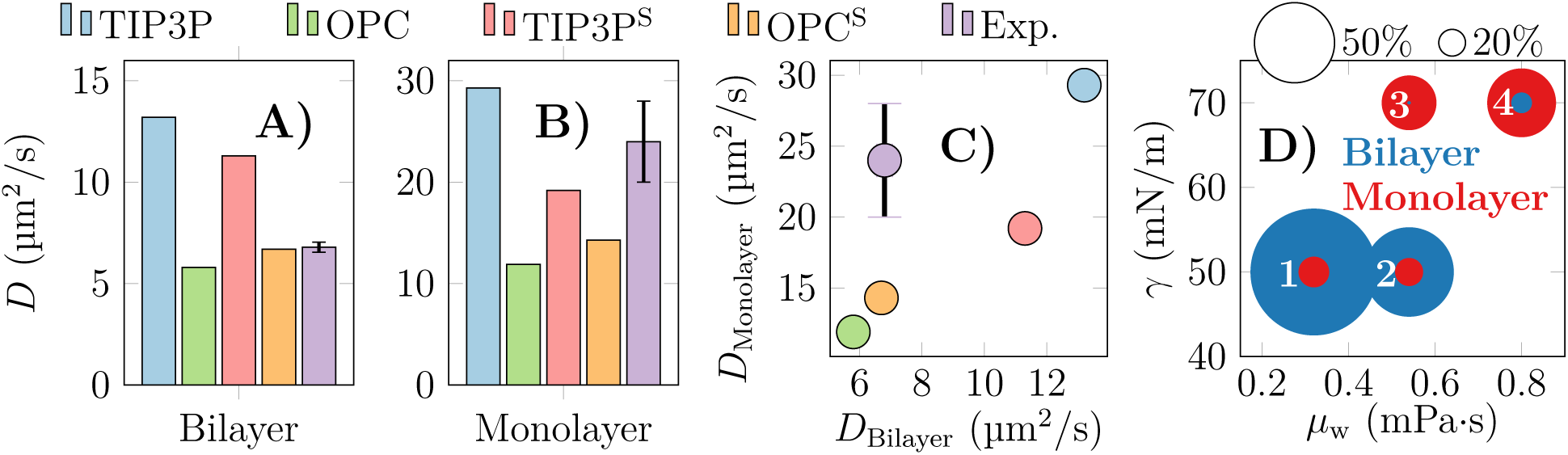
Diffusion coefficients of POPC with different water models and their deviation from experiments. A & B) PBC-corrected data for the POPC bilayer (A) and monolayer (B) are compared to experiments. The used models are the CHARMM-specific TIP3P, OPC, and their mass-scaled versions that have mutually equal shear viscosities. It is noteworthy that the blue and red bars as well as the green and yellow bars (*i.e.* the regular and mass-scaled versions of the water models) should have the same thermodynamic properties yet different dynamic properties, whereas the red and yellow ones should have the same dynamic yet different thermodynamic properties. C) The diffusion coefficients represented on a 2D map to simultaneously highlight the deviations of the bilayer and monolayer values from experiments. D) Deviation (in %) from the experimental estimates mapped to the shear viscosity and surface tension of the water models. The marker size shows the magnitude of the deviation, and the colours correspond to the monolayers and bilayers. The models are: 1) TIP3P, 2) TIP3P^S^, 3) OPC^S^, and 4) OPC.

Finally, Table 1 also contains the shear viscosities of the bilayers (*µ*_bil_) and monolayers (*µ*_mono_) extracted from simulations, as well as the Saffman–Delbrück lengths (*L*_SD_). With the interleaflet coupling explicitly accounted for, one would again assume that the shear viscosities of the monolayers and bilayers would be similar. However, there is a large variation. In the case of TIP3P water; for SOPC the values are similar; for POPC and OMPC the bilayer shear viscosity is ≈1.5-fold larger; and for DOPC the bilayer is twice as viscous as the monolayer. In the case of OPC water, the bilayer viscosities are always larger by ≈10% (OMPC) or by ≈40% (other lipids). These differences are quite striking, and certainly indicate that an equal APL alone is insufficient to establish dynamic similarity between monolayers and bilayers. Still, the relation between the diffusion coefficients and membrane viscosities seem to follow two trends, one for the monolayers and one for the bilayers across the two studied water models (Fig. S6 in the SI), which could help guide future attempts in refining the lipids models.

## 3 Conclusions

We have evaluated the ability of state-of-the-art atomistic molecular dynamics simulations to describe the lateral dynamics of lipids in bilayers and monolayers. To properly compare our simulation results to experiments, we exploited the latest developments for the extraction of reliable diffusion coefficient estimates^38,50^ and for the elimination of finite size effects.^35,36,39^ We simulated four single-component lipid bilayers and monolayers composed of a set of phosphocholine lipids with varying acyl chain composition, for which diffusion coefficients were measured in our recent study.^48^

The bilayers were studied in their native tensionless state, whereas the monolayers were confined to a matching area per lipid. At this equal surface density, the monolayers and bilayers were structurally similar (Figs. S2 and S3, see also Ref. 48). The bilayer simulations confirmed the earlier observation that the CHARMM36 force field significantly overestimates lipid diffusion coefficients when the finite-size effects are properly accounted for.^32,37^ This issue is easily concealed by the fact that the diffusion coefficients extracted from our small and intermediate simulation systems underestimate the experimental estimates, whereas our large simulation system provides a good agreement with experiment (Fig. S1).

In line with Ref. 36, we found interleaflet friction to have little effect on the corrected diffusion coefficient estimates. Moreover, the parameter values extracted from shearing simulations were in reasonable agreement with experiment, indicating that they are not the source of the too fast diffusive dynamics in CHARMM36. An even more worrying aspect is that even the trends in the magnitudes of the diffusion coefficients were not reproduced by the simulations.

To our knowledge, our study represents the first attempt to correct for finite size effects in lipid monolayers. In the finite size correction, we treated monolayers using the same parameters as the bilayers except that the interleaflet friction was set to zero. Using the CHARMM-specific TIP3P water model recommended for CHARMM36, we observed an excellent agreement between our simulations and experiments (Fig. 3).

The change of water models from TIP3P into OPC with a significantly more accurate representation of shear viscosity resulted in a decrease of the bilayer diffusion coefficients and a good agreement with experiment. For monolayers, the change into OPC also affects the surface tension of the monolayer-covered interface, and the change leads to an underestimation of experimental values by a factor of 2.

Finally, we developed two mass scaled water models based on TIP3P and OPC that had equal shear viscosities. These TIP3P^S^ and OPC^S^ models provided more similar yet still very different diffusion coefficients than the unscaled TIP3P and OPC models (Fig. 4, see also independent values in Table S3), indicating that properties other than viscosity play key roles in determining lipid dynamics. These additional simulations also suggested that reproducing monolayer diffusion coefficients with current lipid models is only feasible using a water model with incorrect shear viscosity, and hence the reparametrization of the lipid models might be required. We also hope that our results highlight the numerous pitfalls of extracting dynamic information from MD simulations.

## 4 Experimental

### 4.1 Simulation Setups

We performed three sets of simulations: equilibrium simulations of lipid bilayers and lipid monolayers and non-equilibrium shearing simulations of lipid bilayers. In all sets, three types of phosphocholine lipids with varying acyl chains were considered: 1,2-dioleoyl-*sn*-glycero-3-phosphocholine (DOPC) with two monounsaturated oleate chains, 1-stearoyl-2-oleoyl-*sn*-glycero-3-phosphocholine (SOPC) with a 18-carbon long saturated stearate chain, 1-palmitoyl-2-oleoyl-*sn*-glycero-3-phosphocholine (POPC) with a 16-carbon long saturated palmitate chain, and 1-oleoyl-2-myristoyl-*sn*-glycero-3-phosphocholine (OMPC) with a 14-carbon long myristate chain. Notably, the acyl chain expected to be in a more extended conformation always occupied the *sn*-1 position; in SOPC and POPC this was the saturated chain and in OMPC the unsaturated oleate chain. For all simulations covering all four lipid types, the CHARMM36 lipid model^45^ and either the CHARMM-specific TIP3P water model^46,47^ or the 4-point OPC^54^ were used. For the non-equilibrium simulations, only TIP3P was considered. Additionally, POPC monolayer and bilayer simulations were repeated with mass-scaled TIP3P^S^ and OPC^S^ models. While other lipids are readily available in CHARMM, we generated the OMPC topology by ourselves. All the topologies are available in the respective Zenodo uploads.

For the equilibrium simulations, all systems were simulated in three sizes with a total of 64, 256, and 1024 lipids. Each system was hydrated by 50 water molecules per lipid. The smallest bilayers were set up using CHARMM-GUI and subjected to the CHARMM equilibration protocol.^60^ After that, the atomic coordinates were replicated to create 4- (2 × 2) or 16-fold (4 × 4) larger systems. All generated bilayers were simulated for 100 ns using the recommended settings (see details below). The bilayer APLs stabilized during these simulations, after which the bilayers were transformed into corresponding monolayers setups in which the two monolayers were separated by a water slab. This was achieved by translating the coordinates of one of the leaflets while simultaneously adding a large vacuum separating the acyl chains across the periodic boundary conditions, preserving the APL. Eventually, all monolayers and bilayers using TIP3P water were simulated using GROMACS 2021.3, whereas those with OPC water used GROMACS 2023.3.^44^ All simulations were 1 µs long, and the first 100 ns were omitted from all analyses.

For the non-equilibrium simulations, we extended the thickness of the water slab by 17.5 nm, resulting in a water-to-lipid ratio of ≈230. We performed the shearing simulations using a tool developed to study the unfolding of proteins under water shear. ^61^ As the program takes in the *x* coordinate intervals in which force is applied (to water molecules), we rotated the system so that the membrane was aligned with the (*y, z*) plane. Opposite forces of equal magnitude were applied to the water molecules in the *x* intervals of [0,4] nm and [max(*x*)-4, max(*x*)] nm. The applied force constants were 100, 200, 300, 400, and 500 kJ/(mol·nm) and they were used without a weighting profile. These force constants correspond to ≈170– 830 pN forces. Considering that there are ≈10,900 water molecules in the slab onto which this force is applied, the average force per molecule is ≈0.015–0.076 pN. These forces resulted in velocity gradients similar to those in a recent study which used a different code to induce them.^39^

We computed the membrane dimensions as in our earlier work, ^32,37^ namely by using the final simulation box dimensions for box edge and height that are required for the PBC correction. The membrane thickness was extracted by fitting Gaussian peaks to the phosphorus atom density profiles along the membrane normal.

For the shear viscosity of the CHARMM-specific TIP3P model at 298 K, we use the value previously calculated by Ong and Liow ^57^. The sheared bilayers contained a total of 256 lipid molecules, hence representing the medium system size in our study. The largest bilayer was omitted as the shearing simulation scaled poorly with an increasing number of water molecules onto which the shearing force was applied.

### 4.2 Simulation Parameters

For all simulations, we used the leap-frog integrator with a time step of 2 fs. Buffered Verlet lists^62^ were used to keep track of atomic neighbors. The Lennard-Jones potential was cut of at 1.2 nm, and the forces were switched to zero between 1.0 and 1.2 nm. The smooth Particle Mesh Ewald algorithm^63^ was used to calculate long-range contributions to electrostatics.

The non-equilibrium shear simulations and the monolayer simulations were performed in the NVT ensemble, whereas NPT was used for the bilayers. The temperature of the membrane and the solvent were separately maintained at 298.15 K using the stochastic velocity rescaling thermostat^64^ with a time constant of 1 ps. The bonds involving hydrogens were constrained using P-LINCS.^65^ For bilayers, the Parrinello–Rahman barostat^66^ with a target pressure of 1 bar, time constant of 5 ps, and compressibility of 4.5×10^−5^ bar^−1^ was used. The two dimensions (*x, y*) along the membrane plane were coupled together and the dimension normal to the membrane (*z*) separately in a semi-isotropic coupling scheme.

### 4.3 Analyses

The three system sizes allowed for the elimination of PBC effects and the extraction of *D_∞_*, which can be directly compared with experiments. To this end, we calculated the *D*_PBC_ from the three different system sizes and fitted these values with the full numerical solution for a monotopic inclusion and involving finite interleaflet friction (*b*). The values of *b* were extracted from non-equilibrium simulations as explained below. The fitting formula, evaluated numerically, is too tedious to include here, so we refer the reader to the Supplemental Material of Ref. 36.

We also used the same numerical framework for the monolayers. Here, the thickness of the solvent layer separating the monolayers is equal to that in the bilayers, and hence the same *L_z_* value was used for both. On the acyl chain side, the monolayers are separated by a thick vacuum slab (Fig. 1C), and hence we modelled this as a vanishing interleaflet friction (*b* = 0) in the fits. For all systems, the *D*_PBC_ values were obtained from mean squared displacement using the framework developed recently.^38,50^

For the sake of completeness, we also compared the results from the full numerical solution for monotopic inclusions to those obtained assuming a bitopic inclusion. Finally, we also used an approximate formula which Vögele et al. found to provide a reasonable agreement for membrane lipids. This latter reads

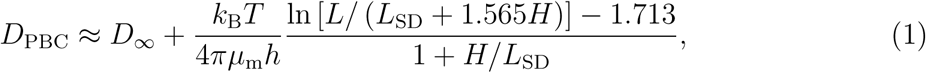

where *D*_PBC_ is the value extracted from a simulation performed using PBCs and *D_∞_* is a value with the PBC effects eliminated. *k*_B_ is the Boltzmann constant, *T* the temperature, *h* membrane thickness, *µ*_m_ the shear viscosity of the membrane, *H* half the thickness of the solvent layer in the simulation, and *L*_SD_ = *µ*_m_*h/*(2*µ*_w_) is the SD length with *µ*_w_ the shear viscosity of the solvent (water).

For the non-equilibrium simulations, we followed the approach of Zgorski et al..^39^ Hence, we applied forces to the water molecules located far from the bilayer. These forces, which had opposing direction on the two sides of the bilayer, resulted in a linear velocity gradient and hence a shear rate in the vicinity of the bilayer (Fig. S10A). This total shear rate summed over the two sides of the membrane *ζ̇*_total_ = *ζ̇*_upper_ + *ζ̇*_lower_ corresponds to a shear force *F* = *µ*_w_*ζ̇*_total_. The shear rate *ζ̇*_total_ induces the relative displacement of the two bilayer leaflets at constant velocity *v* (Fig. S10B). The shear force and the displacement velocity are related by *F* = *bv*, and hence we can reorganize to obtain ^39^

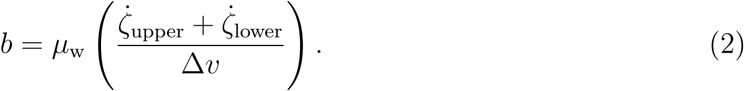

For the TIP3P, a value for *µ*_w_ of 0.322 mPa·s was used.^57^ The values of *b* were measured at five different applied forces *f* , *i.e.* five different total shear rates *ζ̇*_total_. The *b* values were independent of the applied force (Fig. S10C), and the reported values are mean ± standard deviation of the five values.

The mass-scaled water models were evaluated in terms of their shear viscosity and surface tension. The viscosity simulations were carried out in the NVT ensemble with 832 water molecules, a timestep of 2 fs, and a total sampling time of 10 ns, saving the pressure results every 6 fs (3 timesteps). The Green–Kubo formula,

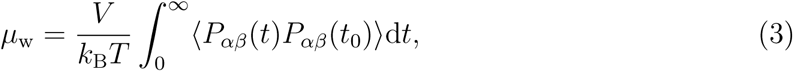

was used to extract the viscosity.^67^ Here, *k*_B_, *V* , and *T* are the Boltzmann constant, the volume of the simulation box, and the temperature, respectively. We have calculated the surface tension for each mass ratio to ensure that the mass change does not affect the interaction between particles and, hence, the surface tension. For the surface tension calculation, one of the box vectors was increased to 15 nm to include a thick enough vacuum slab to avoid interactions over the periodic boundaries, and the surface tension values were extracted from the pressure tensor (see, *e.g.*, Ref. 49).

## Acknowledgement

M.J. thanks the Research Council of Finland (grant no. 338160) for funding, CSC–IT Center for Science (Espoo, Finland) for computational resources. T.M. and S.C. are grateful to the Postgraduate Scholarship Program of the University of Potsdam.

## Supporting Information Available

**TOC Graphic**

