## Supplementary Material for "Challenges in the Accurate Modelling of Lipid Dynamics in Monolayers and Bilayers"

Table S1: **Diffusion coefficients of individual simulations with the TIP3P water model.** The diffusion coefficient values (in $10^{-8}$ cm$^2$/s) are listed for each lipid type and each system size. The APL (in Å$^2$) and $\gamma$ (in mN/m) are also listed to demonstrate that these properties are consistent between the different system sizes.

| | | **Bilayer** | | **Monolayer** | |
|---|---|---|---|---|---|
| Lipid | Lipids | APL | $D_{\mathrm{bil}}^{\mathrm{PBC}}$ | $\gamma$ | $D_{\mathrm{mono}}^{\mathrm{PBC}}$ |
| OMPC | 64 | $64.7 \pm 1.2$ | $4.6 \pm 0.9$ | 23.0 | $6.7 \pm 1.4$ |
| OMPC | 256 | $64.5 \pm 0.8$ | $6.2 \pm 1.3$ | 28.0 | $11.8 \pm 2.3$ |
| OMPC | 1024 | $64.4 \pm 0.3$ | $7.7 \pm 1.5$ | 25.6 | $13.6 \pm 2.6$ |
| DOPC | 64 | $67.7 \pm 1.0$ | $5.0 \pm 1.0$ | 28.3 | $6.9 \pm 1.4$ |
| DOPC | 256 | $67.7 \pm 1.1$ | $6.2 \pm 1.2$ | 30.2 | $10.7 \pm 2.1$ |
| DOPC | 1024 | $67.6 \pm 0.4$ | $7.2 \pm 1.5$ | 31.9 | $14.0 \pm 2.8$ |
| POPC | 64 | $63.5 \pm 1.3$ | $3.8 \pm 0.8$ | 31.4 | $6.7 \pm 1.3$ |
| POPC | 256 | $63.8 \pm 0.6$ | $5.5 \pm 1.1$ | 33.4 | $11.0 \pm 2.1$ |
| POPC | 1024 | $63.7 \pm 0.4$ | $6.4 \pm 1.3$ | 30.6 | $11.8 \pm 2.3$ |
| SOPC | 64 | $63.4 \pm 1.8$ | $3.1 \pm 0.7$ | 32.4 | $6.3 \pm 1.3$ |
| SOPC | 256 | $63.4 \pm 0.8$ | $4.8 \pm 1.0$ | 29.9 | $7.6 \pm 1.5$ |
| SOPC | 1024 | $63.3 \pm 0.4$ | $5.6 \pm 1.1$ | 32.7 | $10.6 \pm 2.1$ |

Table S2: **Diffusion coefficients of individual simulations with the OPC water model.** The diffusion coefficient values (in $10^{-8}$ cm$^2$/s) are listed for each lipid type and each system size. The APL (in Å$^2$) and $\gamma$ (in mN/m) are also listed to demonstrate that these properties are consistent between the different system sizes.

| | | **Bilayer** | | **Monolayer** | |
|---|---|---|---|---|---|
| Lipid | Lipids | APL | $D_{\mathrm{bil}}^{\mathrm{PBC}}$ | $\gamma$ | $D_{\mathrm{mono}}^{\mathrm{PBC}}$ |
| OMPC | 64 | $63.4 \pm 0.2$ | $2.0 \pm 0.4$ | 33.4 | $3.5 \pm 0.7$ |
| OMPC | 256 | $63.3 \pm 0.1$ | $2.9 \pm 0.6$ | 26.0 | $4.4 \pm 0.9$ |
| OMPC | 1024 | $61.1 \pm 0.0$ | $3.7 \pm 0.8$ | 25.5 | $5.5 \pm 1.2$ |
| DOPC | 64 | $65.9 \pm 0.1$ | $2.2 \pm 0.5$ | 27.9 | $3.0 \pm 0.6$ |
| DOPC | 256 | $66.1 \pm 0.1$ | $3.0 \pm 0.6$ | 30.8 | $4.5 \pm 0.9$ |
| DOPC | 1024 | $66.0 \pm 0.0$ | $3.5 \pm 0.7$ | 31.0 | $5.8 \pm 1.2$ |
| POPC | 64 | $62.3 \pm 0.2$ | $1.9 \pm 0.4$ | 27.9 | $2.4 \pm 0.5$ |
| POPC | 256 | $62.6 \pm 0.1$ | $2.6 \pm 0.5$ | 30.7 | $3.9 \pm 0.8$ |
| POPC | 1024 | $62.4 \pm 0.0$ | $3.1 \pm 0.7$ | 29.2 | $4.6 \pm 1.0$ |
| SOPC | 64 | $62.4 \pm 0.3$ | $1.7 \pm 0.4$ | 28.1 | $2.1 \pm 0.5$ |
| SOPC | 256 | $62.1 \pm 0.1$ | $2.4 \pm 0.5$ | 30.2 | $3.1 \pm 0.7$ |
| SOPC | 1024 | $62.1 \pm 0.0$ | $2.8 \pm 0.6$ | 30.2 | $4.3 \pm 0.9$ |

Table S3: **Diffusion coefficients of individual simulations with the mass-scaled water models.** The diffusion coefficient values (in $10^{-8}$ cm$^2$/s) are listed for POPC wiht the two modified water models based on TIP3P and OPC. The APL (in Å$^2$) and $\gamma$ (in mN/m) are also listed to demonstrate that these properties are consistent between the different system sizes.

| | | **Bilayer** | | **Monolayer** | |
|---|---|---|---|---|---|
| Lipid | Lipids | APL | $D_{\mathrm{bil}}^{\mathrm{PBC}}$ | $\gamma$ | $D_{\mathrm{mono}}^{\mathrm{PBC}}$ |
| **TIP3P$^{\mathbf{S}}$** | | | | | |
| POPC | 64 | $64.0 \pm 0.2$ | $3.3 \pm 0.7$ | 26.0 | $6.3 \pm 1.3$ |
| POPC | 256 | $64.0 \pm 0.1$ | $4.7 \pm 1.0$ | 26.0 | $6.2 \pm 1.3$ |
| POPC | 1024 | $63.9 \pm 0.1$ | $5.7 \pm 1.2$ | 31.4 | $10.6 \pm 2.1$ |
| **OPC$^{\mathbf{S}}$** | | | | | |
| POPC | 64 | $62.5 \pm 0.1$ | $2.2 \pm 0.5$ | 30.1 | $3.1 \pm 0.7$ |
| POPC | 256 | $62.5 \pm 0.1$ | $2.9 \pm 0.6$ | 30.7 | $4.5 \pm 0.9$ |
| POPC | 1024 | $62.4 \pm 0.0$ | $3.5 \pm 0.7$ | 32.6 | $5.8 \pm 1.2$ |

Table S4: **Parameters extracted from the shearing simulations.** The result is shown for the four lipids and with 5 shearing rates for each. See the Methods section in the main text for the descriptions of these parameters.

| $F$ (kJ/(mol·nm)) | $\dot{\gamma}_{\text{upper}}$ (ns$^{-1}$) | $\dot{\gamma}_{\text{lower}}$ (ns$^{-1}$) | $v$ ($10^{-3}$ m/s) | $b$ ($10^{7}$ Pa·s/m) |
|---|---|---|---|---|
| | | **DOPC** | | |
| 100 | -1.098 $\pm$ 0.003 | -1.093 $\pm$ 0.004 | 15.3 $\pm$ 0.1 | 46.02 $\pm$ 0.15 |
| 200 | -2.240 $\pm$ 0.003 | -2.231 $\pm$ 0.004 | 30.2 $\pm$ 0.2 | 47.67 $\pm$ 0.07 |
| 300 | -3.306 $\pm$ 0.005 | -3.257 $\pm$ 0.005 | 43.1 $\pm$ 0.1 | 49.03 $\pm$ 0.07 |
| 400 | -4.294 $\pm$ 0.010 | -4.262 $\pm$ 0.010 | 59.2 $\pm$ 0.1 | 46.54 $\pm$ 0.11 |
| 500 | -5.204 $\pm$ 0.020 | -5.204 $\pm$ 0.011 | 76.5 $\pm$ 0.1 | 43.81 $\pm$ 0.13 |
| | | **SOPC** | | |
| 100 | -1.229 $\pm$ 0.003 | -1.151 $\pm$ 0.003 | 23.8 $\pm$ 0.1 | 32.25 $\pm$ 0.08 |
| 200 | -2.446 $\pm$ 0.004 | -2.275 $\pm$ 0.004 | 54.0 $\pm$ 0.1 | 28.14 $\pm$ 0.05 |
| 300 | -3.543 $\pm$ 0.007 | -3.351 $\pm$ 0.006 | 54.7 $\pm$ 0.1 | 40.62 $\pm$ 0.08 |
| 400 | -4.561 $\pm$ 0.020 | -4.267 $\pm$ 0.010 | 81.0 $\pm$ 0.1 | 35.10 $\pm$ 0.12 |
| 500 | -5.600 $\pm$ 0.020 | -5.091 $\pm$ 0.020 | 111.3 $\pm$ 0.2 | 30.93 $\pm$ 0.12 |
| | | **POPC** | | |
| 100 | -1.247 $\pm$ 0.003 | -1.178 $\pm$ 0.002 | 22.6 $\pm$ 0.2 | 34.57 $\pm$ 0.07 |
| 200 | -2.450 $\pm$ 0.004 | -2.271 $\pm$ 0.003 | 46.9 $\pm$ 0.1 | 32.41 $\pm$ 0.05 |
| 300 | -3.557 $\pm$ 0.007 | -3.290 $\pm$ 0.004 | 72.2 $\pm$ 0.1 | 30.52 $\pm$ 0.05 |
| 400 | -4.630 $\pm$ 0.010 | -4.280 $\pm$ 0.010 | 108.6 $\pm$ 0.1 | 26.42 $\pm$ 0.06 |
| 500 | -5.480 $\pm$ 0.020 | -5.167 $\pm$ 0.020 | 128.4 $\pm$ 0.2 | 26.70 $\pm$ 0.10 |
| | | **OMPC** | | |
| 100 | -1.245 $\pm$ 0.002 | -1.189 $\pm$ 0.003 | 21.7 $\pm$ 0.2 | 36.15 $\pm$ 0.07 |
| 200 | -2.444 $\pm$ 0.004 | -2.304 $\pm$ 0.003 | 47.0 $\pm$ 0.1 | 32.50 $\pm$ 0.05 |
| 300 | -3.526 $\pm$ 0.009 | -3.319 $\pm$ 0.008 | 65.9 $\pm$ 0.1 | 33.47 $\pm$ 0.08 |
| 400 | -4.524 $\pm$ 0.020 | -4.298 $\pm$ 0.010 | 94.9 $\pm$ 0.1 | 29.94 $\pm$ 0.10 |
| 500 | -5.541 $\pm$ 0.020 | -5.098 $\pm$ 0.020 | 116.7 $\pm$ 0.1 | 29.36 $\pm$ 0.11 |

Table S5: Diffusion coefficients bilayer lipids simulated with TIP3P water. The values are obtained either 1) with the numerical approach[S1] for monotopic inclusions ($D^{b}_{\text{bil}}$); 2) with the numerical approach[S1] for bitopic inclusions ($D^{\infty}_{\text{bil}}$); or 3) using the analytical expression (Eq. (1) in the main text) and assuming bitopic inclusions ($D^{\text{approx.}}_{\text{bil}}$). Values in $10^{-8}$ cm$^2$/s.

| Lipid | $D^{b}_{\text{bil}}$ | $D^{\infty}_{\text{bil}}$ | $D^{\text{approx.}}_{\text{bil}}$ |
|---|---|---|---|
| OMPC | 15.4 | 16.9 | 17.4 |
| DOPC | 13.1 | 14.1 | 14.6 |
| POPC | 13.2 | 14.7 | 15.2 |
| SOPC | 12.3 | 13.6 | 14.1 |

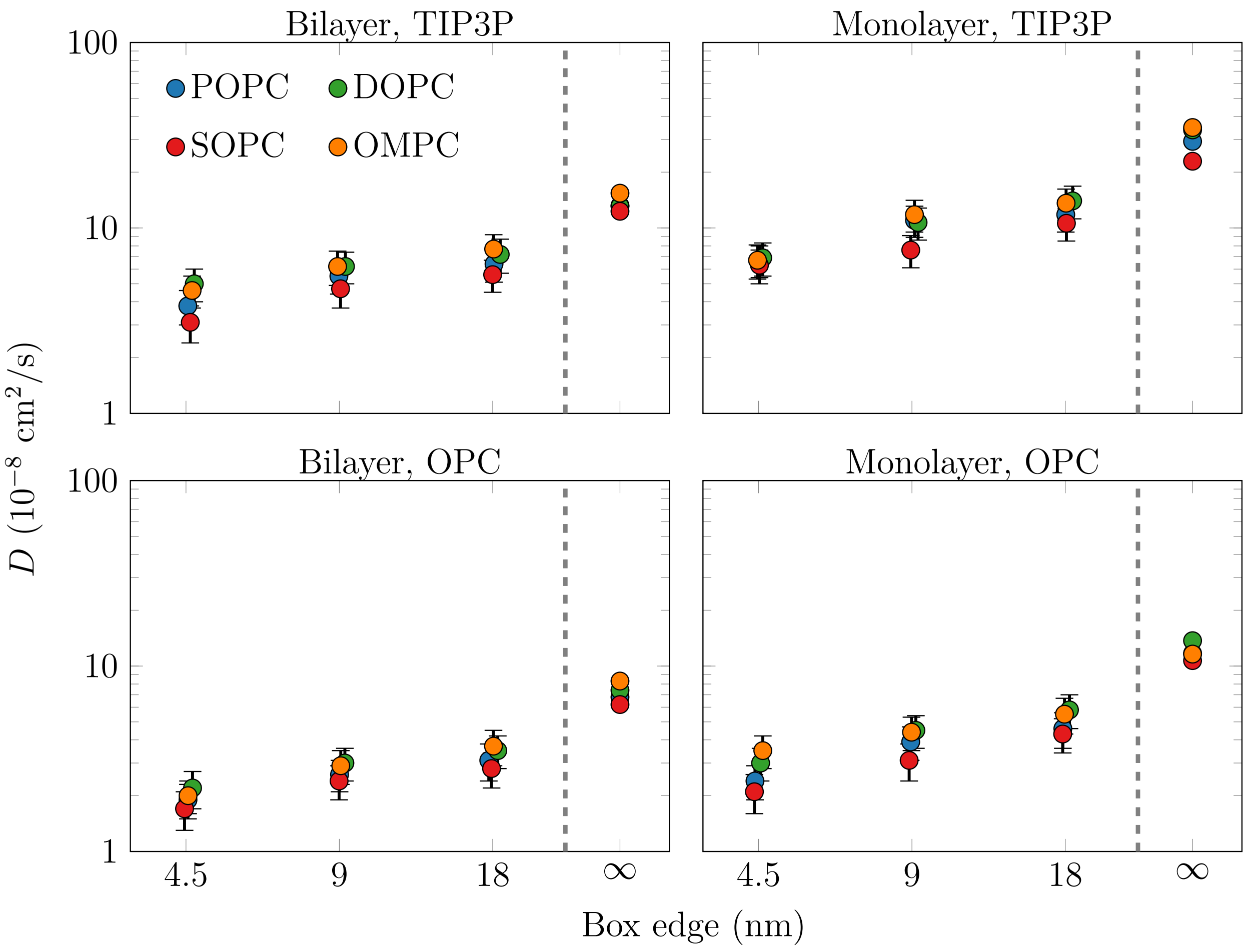


Figure S1: **Size-dependence of diffusion coefficients.** Diffusion coefficient values are shown with markers for the small (box edge≈4.5 nm)), medium (≈9 nm), and large (≈18 nm) system sizes. The markers at ∞ show the values extrapolated to infinite system size, *i.e.* with the PBC effects eliminated. Data are shown for both the bilayers and monolayers and for the TIP3P and OPC water models. Note the logarithmic scale on both axes.

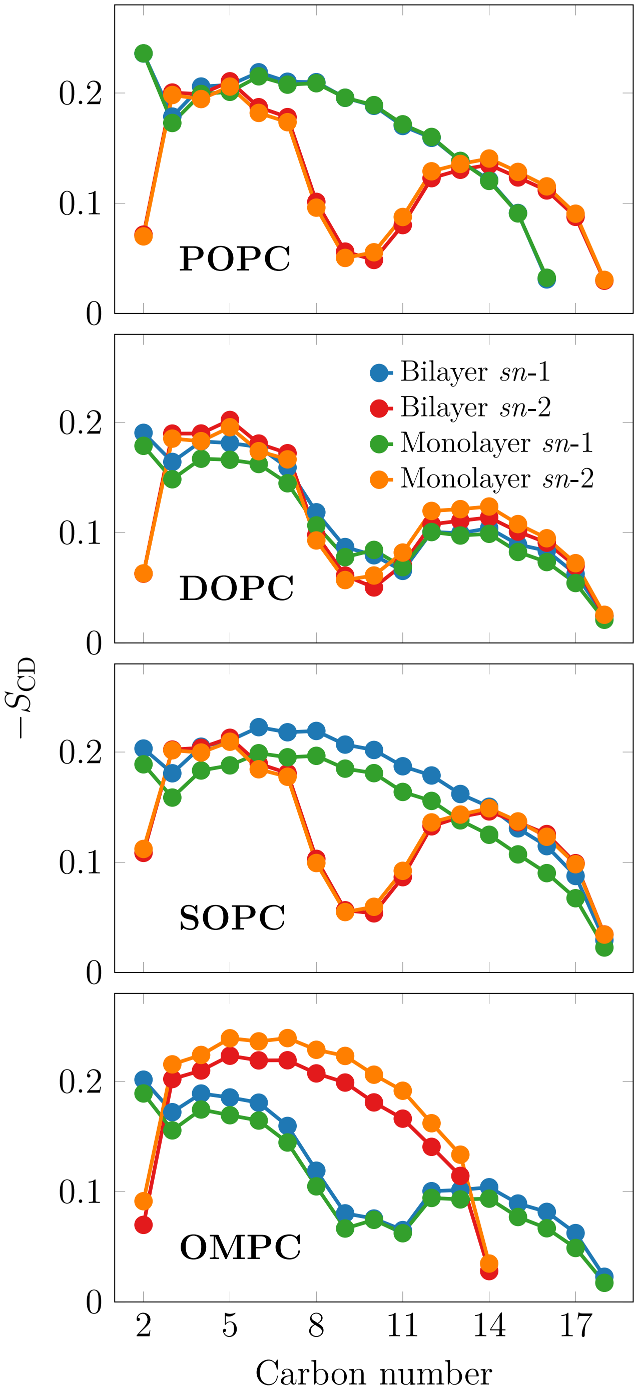


Figure S2: Deuterium order parameters of the acyl chains from simulations with TIP3P. The structure of the corresponding monolayers and bilayers are very similar.

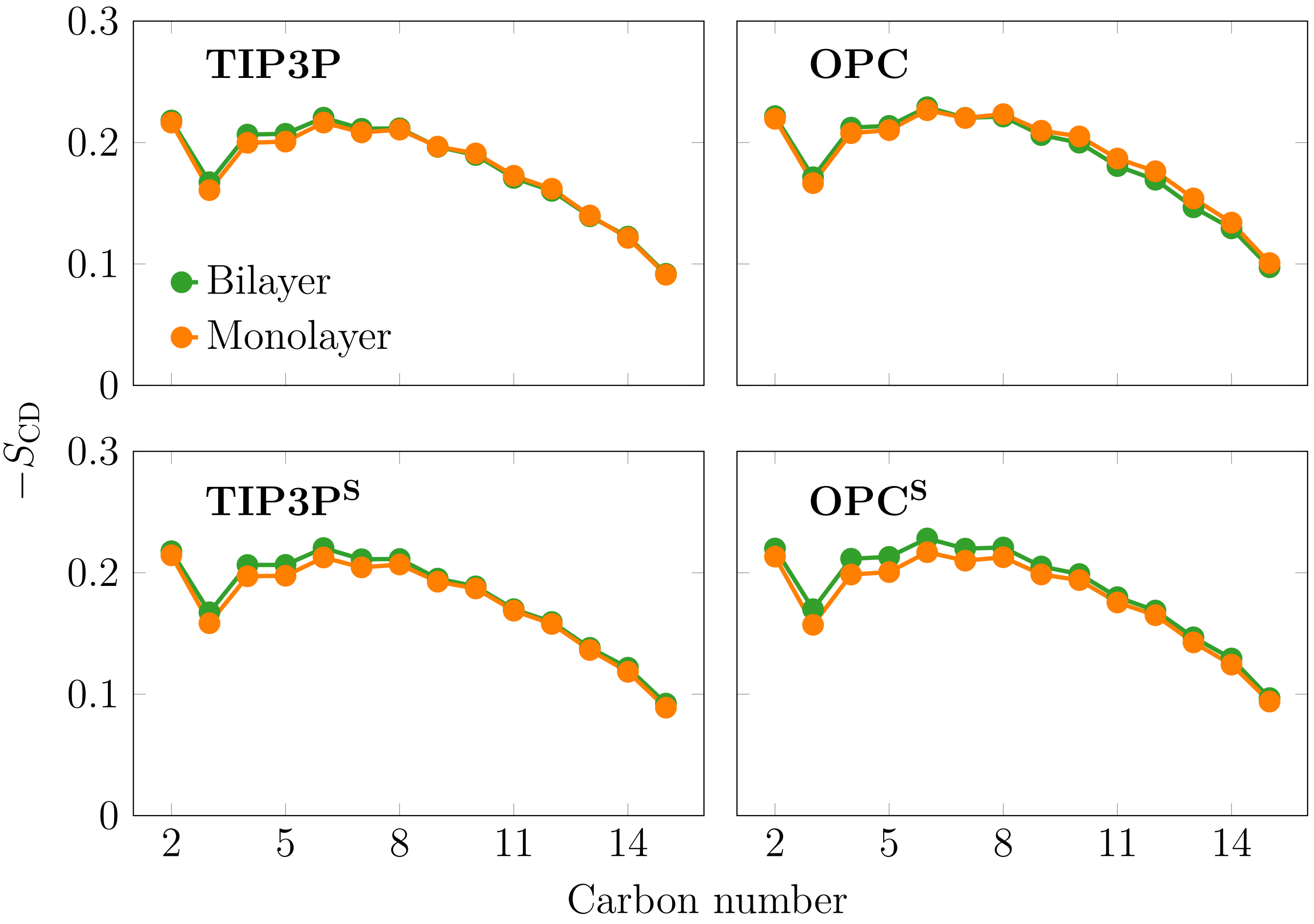


Figure S3: Deuterium order parameters of the *sn*-1 chain of POPC with different water models and in both monolayers and bilayers. Overall, the profiles are very similar across all studied systems.

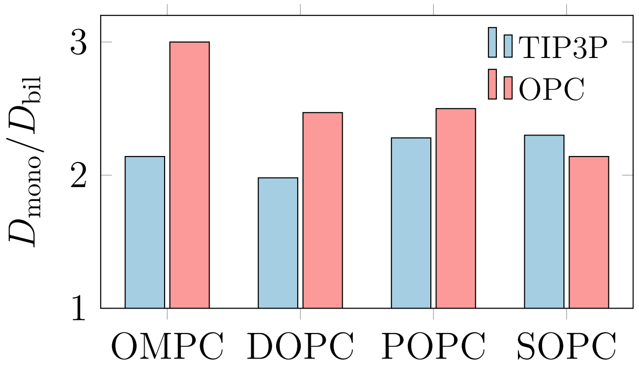


Figure S4: **Ratios of the monolayer and bilayer diffusion coefficients.** The values shown here are PBC-corrected, as explained in the main text.

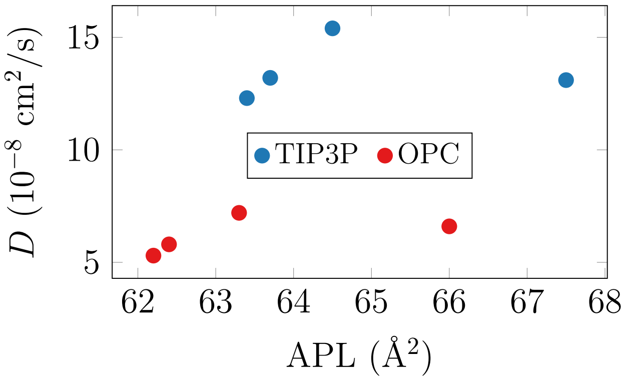


Figure S5: **Effect of the area per lipid on the diffusion coefficients of the lipids.** PBC-corrected data are shown for the bilayer simulations with the TIP3P and OPC water models. No simple free area theory-like dependence ($D \sim \sqrt{A} \times \exp\left(-a_0/(A - a_0)\right)$) with $A$ being the APL is observed, indicating that the lipid chemistry also plays a role.

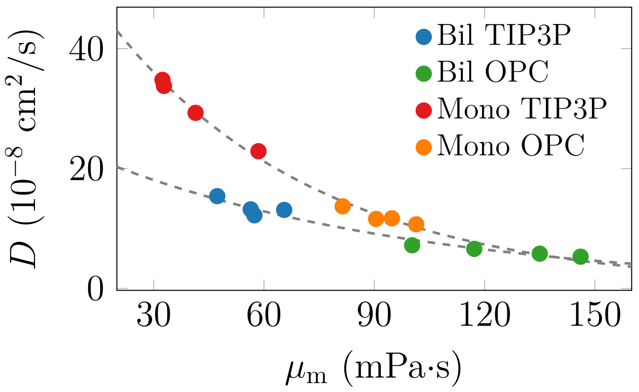


Figure S6: **The relationship between shear viscosity and diffusion coefficient.** Data are shown for the lipid monolayers and bilayers with the TIP3P and OPC water models. Both types of systems follow the same trend (dashed line to guide the eye) for the two used water models.

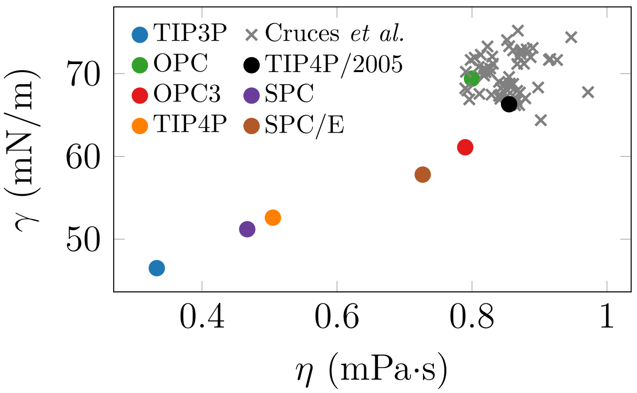


Figure S7: **Correlation of shear viscosity and surface tension of popular water models.** Water models included here are TIP3P and TIP4P,[S2] TIP4P/2005,[S3] SPC[S4] and SPC/E,[S5] OPC,[S6] and OPC3.[S7] The viscosity and surface tension values are taken from Refs. S3,S8–S11. In addition, data for the multiple water models parameterized recently based on density, self-diffusion coefficient, first peak in the oxygen–oxygen radial distribution function, and the dielectric constant, are shown ("Cruces *et al.*") (Private communication; these models were generated using the tools used in Ref. S12, although not reported in the publication).

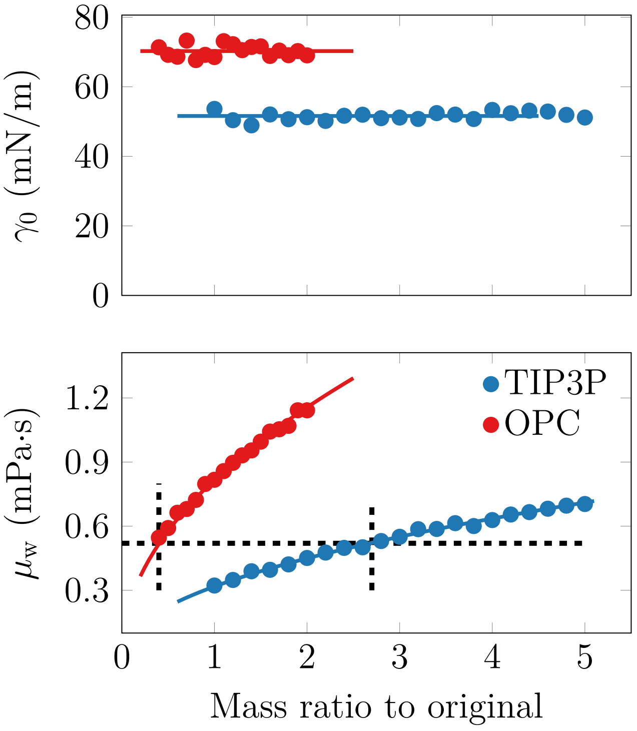


Figure S8: **Scaling of surface tension and shear viscosity with water mass.** Surface tension is unaffected by mass, whereas shear viscosity shows the expected dependence of $\eta_w \sim \sqrt{M}$ with mass $M$. The gray lines highlight the similar viscosities of ≈0.53 mPa·s of the models, which are obtained after scaling the OPC and TIP3P masses by factors of 0.4 and 2.9, respectively. These models are labeled $\mathrm{OPC_S}$ and $\mathrm{TIP3P^S}$ in the main text.

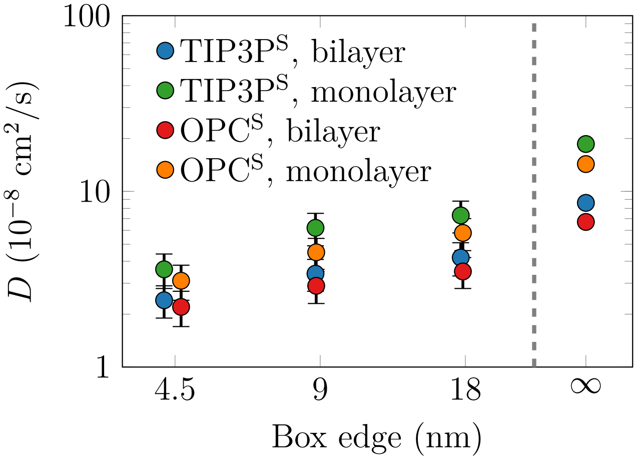


Figure S9: **Size-dependence of diffusion coefficients with the mass-scaled water models.** Diffusion coefficient values are shown with markers for the small (box edge≈4.5 nm)), medium (≈9 nm), and large (≈18 nm) system sizes. The markers at $\infty$ show the values extrapolated to infinite system size, *i.e.* with the PBC effects eliminated. Data are shown for the POPC bilayer and monolayer and for the TIP3P$^S$ and OPC$^S$ water models. Note the logarithmic scale on both axes.

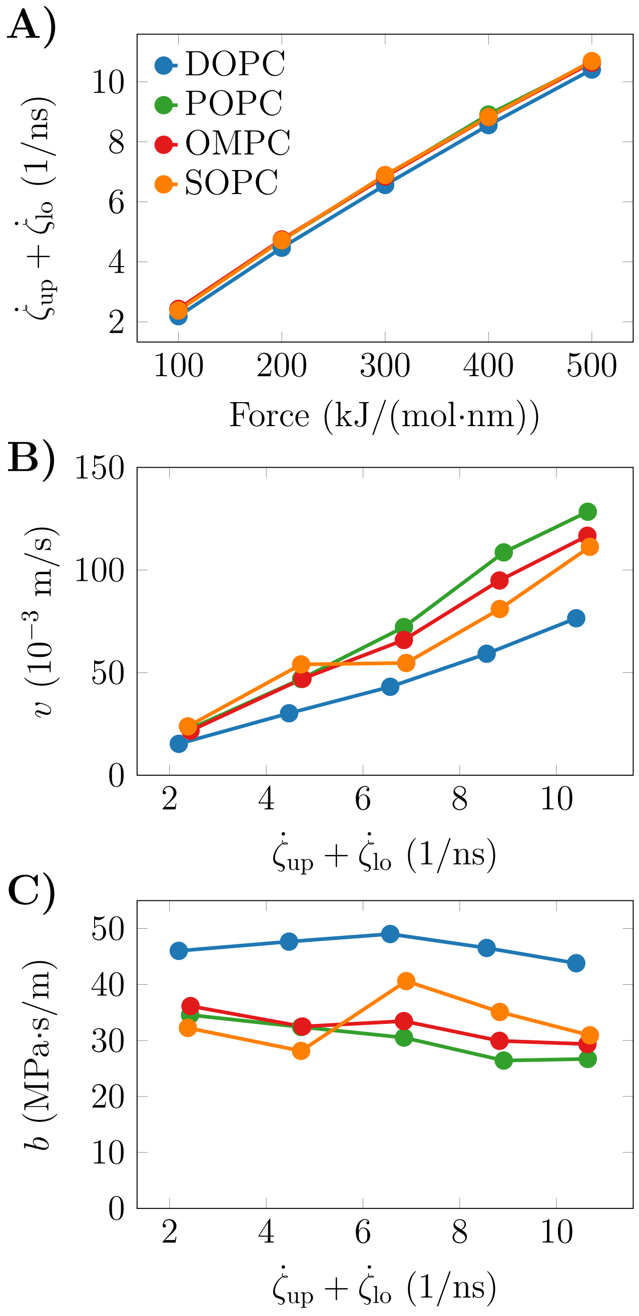


Figure S10: **Results from the shearing simulations. A)** The total shear rate $\dot{\gamma}_{up} + \dot{\gamma}_{lo}$ (sum of shear rates affecting the upper and lower bilayer leaflets) as a function of the force applied to the water slab distant from the bilayer. The shear rate grows linearly as a function of the applied force and no cavities were observed, indicating a well-behaving solvent.[S13] **B)** The relative velocity $v$ of the bilayer leaflets as a function of total solvent shear rate. The trend is linear, justifying the approach.[S14] **C)** The interleaflet friction $b$ values as a function of the total solvent shear rate. The values are independent of the shear rate, as required for reliable estimates of the $b$.[S14]
